# Prior contact lens wear reduces susceptibility of the superficially-injured cornea to bacterial adhesion

**DOI:** 10.1101/2025.08.27.672703

**Authors:** Yujia Yang, Orneika Flandrin, Sara Abboud, Eric Jedel, David J. Evans, Suzanne M. J. Fleiszig

## Abstract

Contact lens wear in humans and mice is consistently associated with asymptomatic corneal parainflammation. Here, we tested the hypothesis that the corneal response to lens wear alone can function to protect it against bacterial adhesion enabled by superficial injury. One eye only of mT/mG-LysMcre mice (cell membranes red; Lyz2^+^ cells green) wore a contact lens for 4-6 days. After lens removal, mice were anesthetized and both corneas superficially-injured before bacterial inoculation with either a mouse eyelid commensal (*Macrococcus epidermidis*) or a corneal pathogen (*Pseudomonas aeruginosa*). Inoculation was repeated hourly for 4 hours under anesthesia before euthanasia. Enucleated eyes were fixed overnight, and adherent bacteria visualized using a universal 16S rRNA-targeted FISH probe (*M. epidermidis*) or Blue Fluorescent Protein (*P. aeruginosa*). Confocal imaging and Imaris software were used to quantify bacterial adhesion and location in the epithelium, and also the number, location and morphology of Lyz2^+^ cells. For both commensal and *P. aeruginosa*, prior lens wear resulted in reduced adhesion to the superficially injured corneas (∼ 55% and ∼ 45% respectively). In both instances this correlated with increased numbers of corneal Lyz2^+^ cells. Other details differed for the two bacterial types. For the commensal, prior lens wear resulted in bacteria penetrating deeper into the epithelium versus contralateral eyes, with Lyz2^+^ cells extending their processes further into the epithelium and localizing closer to the cornea surface. For *P. aeruginosa*, prior lens wear resulted in adherent bacteria closer to the cornea surface, and Lyz2^+^ cells moving further away from it. Moreover, while overall Lyz2^+^ cell sphericity increased for *P. aeruginosa* with prior lens wear versus contralateral eyes, it showed no overall change for the commensal. Lyz2^+^ cell volume in the central cornea was decreased for *P. aeruginosa* with prior lens wear but increased for the commensal. Thus, corneal responses to prior lens wear can quantitively reduce bacterial adhesion to superficially-injured corneas correlating with a Lyz2^+^ cell response for both a commensal and a pathogen, with differences in details for the two bacterial types. How continued lens wear supersedes this protective response to promote *P. aeruginosa* infection pathogenesis remains to be determined, as does the relationship to the Lyz2^+^ cell response.

**Author summary:** Contact lens wear induces asymptomatic corneal parainflammation in humans and mice involving multiple immune cell types. Here, we used a mouse model to test if previous lens wear would subsequently protect superficially-injured corneas against bacterial adhesion by either a murine commensal (*Macrococcus epidermidis*) or an opportunistic pathogen (*Pseudomonas aeruginosa*). Confocal imaging was used to quantify bacterial adhesion/location in addition to Lyz2^+^ immune cell numbers, location and morphology. Prior lens wear for 4-6 days, known to trigger corneal parainflammation, reduced the propensity for bacteria to adhere to corneas that were superficially-injured after the lens was removed: ∼ 55% for the commensal and ∼ 45% for *P. aeruginosa* versus contralateral eyes that had not worn a lens. In both instances this correlated with increased Lyz2^+^ immune cell numbers. However, differences were also noted, including that adherent commensals were found penetrating further into the epithelium while adherent *P. aeruginosa* remained close to the corneal surface. Lyz2^+^ cell location, sphericity and volume also differed for the two bacterial types. In sum, lens wear in the absence of bacterial inoculation can enable a response that protects the cornea against bacterial challenge after lens removal. How lens wear *during* inoculation overcomes this protective response to promote *P. aeruginosa* infection pathogenesis remains to be determined.

## Introduction

Contact lenses are widely used for vision correction and therapeutic management of numerous eye pathologies (1,2), yet their wear can lead to adverse events, the most serious being sight-threatening corneal infection (3,4). It is important to understand the mechanisms by which contact lens wear can predispose to infection and other adverse events. In addition to human studies, researchers have developed various lens wearing animal models to study the impact of lens wear on the cornea and ocular surface and to develop lenses with therapeutic value (5–10).

More recently our laboratory developed a murine model of lens wear using lenses custom-designed to fit the mouse eye (11). Using this model, we showed that mice wearing lenses inoculated with *Pseudomonas aeruginosa*, a leading cause of contact lens-related corneal infection (12,13), developed infectious pathology from 1-13 days after lens insertion (11). Interestingly, in the absence of bacterial inoculation, lens wear was associated with subclinical inflammatory responses involving increased numbers of CD11c^+^ cells (after 24 h), Lyz2^+^ cells (at 7 days) and Ly6G^+^ cells (5-6 days) (11). Subsequent studies showed that 24 h of lens wear also involved increased numbers of MHC-II^+^ cells and γδ T cells (14) and that 6 day Ly6G^+^ cell responses required γδ T cells and IL-17A (15). Subclinical corneal inflammation induced by 24 h of lens wear was shown to only resolve at 7 days after lens removal (16). These lens induced immune cell changes in mice resembled changes in Langerhans cell density shown in human corneas as early as 2 h after contact lens insertion (17) and also after 7 days of daily disposable lens wear (18). We termed the subclinical lens-induced corneal inflammation in mice as parainflammatory as postulated by Dr. Nathan Efron for human wear (19). However, it is not yet known if these subclinical corneal responses in mice (or in humans) are beneficial or detrimental to corneal health or if they represent a precursor to clinically evident inflammation or infection.

The purpose of the present study was to use the murine model of lens wear to test the hypothesis that corneal responses to lens wear after 4-6 days would have a protective effect against bacterial adhesion, an important first step in the development of infection. Since the murine cornea is normally remarkably resistant to bacterial adhesion (20,21), we utilized a superficial injury (tissue paper blotting) model to promote bacterial adhesion without subsequent epithelium traversal (22), then compared the impact of prior lens wear versus no lens wear on corneal defense against bacterial adhesion. A murine ocular commensal bacterium (*Macrococcus epidermidis*) and the opportunistic pathogen *P. aeruginosa* were used tested and differences in bacterial adhesion were correlated with quantitative and morphological changes to corneal Lyz2^+^ immune cells.

## Materials and methods

### Ethics statement

All procedures involving mice were carried out in accordance with a protocol (AUP-2019-06-12322) approved by the Animal Care and Use Committee, University of California, Berkeley which is an AAALAC accredited institution. The protocol adheres to PHS policy on the humane care and use of laboratory animals, and the guide for the care and use of laboratory animals. Procedures adhered to the ARVO Statement for the use of Animals in Ophthalmic Vision Research.

### Mice

Six-week-old male and female mT/mG-LysMcre mice were used. These mice were derived from the cross (F1) of a mT/mG mouse (all cell membranes, red) with a LysMcre mouse (Lyz2^+^ cells, myeloid-derived, green) allowing visualization of Lyz2^+^ cells (green) within the cornea (red) (11).

### Contact lens fitting

Mice were lightly anesthetized with 1∼3 % isoflurane using a precision vaporizer (VetEquip Inc., Pleasanton, CA, USA). A custom-made silicone-hydrogel mouse contact lens was fitted onto the right eye of each mouse as previously described (11) while contralateral eyes did not wear a lens. Briefly, after anesthesia, an Elizabethan collar (Kent Scientific, Torrington, CT, USA) was applied to each mouse and contact lenses were fitted using a Handi-Vac suction pen (Edmund Optics, Barrington, NJ, USA). After lens insertion, mice were individually housed with Pure-o’Cel paper bedding (The Andersons Inc., Maumee, OH, USA), and lenses worn for 4 to 6 days.

### Bacterial adhesion assays

Two bacterial species were used for bacterial adhesion assays. A coagulase-negative murine eyelid commensal (*Macrococcus epidermidis*) and the opportunistic pathogen *Pseudomonas aeruginosa* expressing Blue Fluorescent Protein (PAO1F-EBFP2) (23). Bacteria were prepared by growth on a Tryptic Soy Agar (TSA) plate for ∼16 hours at 37 °C then suspended in PBS to a concentration of ∼10^11^ colony-forming units (CFU)/ml. Following contact lens wear, mice were anesthetized by intraperitoneal injection of ketamine (80-100 mg/Kg) and dexmedetomidine (0.25-0.5 mg/Kg). For each mouse both corneas were superficially-injured (blotted) using a Kimwipe ^TM^ tissue paper to promote bacterial adhesion (22) then both inoculated with 5 μl of bacterial suspension once every hour for 4 hours to allow a comparison of bacterial adhesion to the prior lens wear cornea versus the no lens wear cornea. Mice remained anesthetized for the entire 4 hour incubation period and were kept warm on a heated pad. At the end of the incubation period, and before awakening, mice were euthanized by intraperitoneal injection of ketamine (80-100 mg/Kg) and xylazine (5-10 mg/Kg) followed by cervical dislocation. Eyeballs were enucleated, rinsed with PBS, fixed in 2% paraformaldehyde (PFA) overnight at 4°C then processed for quantitative confocal imaging as described below.

### Fluorescence In-situ Hybridization (FISH) labeling

FISH was used to visualize adherent commensal bacteria on whole eyeballs following fixation as previously described (21). Briefly, eyeballs were washed in PBS, 80 % ethanol, and 95 % ethanol sequentially for 10 minutes each at room temperature (RT) followed by incubation in hybridization buffer (0.9 M NaCl, 20 mM Tris-HCl, and 0.01% SDS) at 55°C for 30 minutes. Then a universal 16S rRNA-targeted gene probe [Alexa647]-GCTGCCTCCCGTAGGAGT-[Alexa647] (Eurofins Genomics) was added to the eyeballs at a final concentration of 100 nM before overnight incubation at 55°C. Eyeballs were washed in washing buffer (0.9 M NaCl, 20 mM Tris-HCl) for three times with 10 minutes each at RT and stored in PBS before imaging.

### Imaging and processing

Whole eyeballs were imaged using an Olympus FluoView confocal microscope with a 488 nm laser used for detection of Lyz2^+^-GFP cells and 559 nm laser for the detection of red cell membranes. *P. aeruginosa* expressing BFP was detected using a 405 nm laser. FISH-labeled bacteria were detected using a 635 nm laser. Z stack images were acquired at a 0.8 μm step-size and an aspect ratio of 1024 μm x 1024 μm for bacteria, or at a 1.5 μm step-size and 800 μm x 800 μm aspect ratio for immune cells. Acquired Z stacks were reconstructed as 3-D images using Imaris software (Oxford Instruments, Bitplane AG, Zurich, Switzerland). For adherent bacteria, the ‘Spots’ function was used for quantification. For immune cells, the ‘Surface’ function was used to render the corneal endothelium layer and Lyz2^+^ immune cells. Cell volume, sphericity and distance from endothelium were measured using the Imaris software.

### Statistical analysis

Each group contained 4-6 eyes. A normality test was performed on the accumulated quantitative data which were presented as either the mean ± standard error of the mean (SEM) for parametric data or the median with upper and lower quartiles (median [Q1-Q3]) for non-parametric data as indicated in each figure. For parametric data, a Student’s t-test was applied and for non-parametric data, either a Mann-Whitney U test or Wilcoxon signed-rank test were applied. P < 0.05 was considered significant. Prism 10 was used (GraphPad Software, Boston, MA, USA).

## Results

### Prior contact lens wear reduces commensal bacteria adhesion to the cornea but increases bacterial depth of penetration

We first explored if prior lens wear would impact adhesion of the bacterial commensal to the superficially-injured cornea. Fig. 1 shows significantly fewer bacteria (white) adhered to corneas (red) that had worn a contact lens versus contralateral eyes, a ∼55 % decrease (P = 0.031, Wilcoxon signed-rank test). Despite fewer adherent bacteria, however, prior lens-wearing corneas showed greater depth of bacterial entry into the epithelium. Fig. 2 shows the increased distance of adherent bacteria from the epithelium surface with prior lens wear: NCL 0.83 [0.34 - 1.67] vs. CL 4.71 [2.47 - 8.22] μm (P < 0.0001, Mann-Whitney U test).

**Fig. 1.**
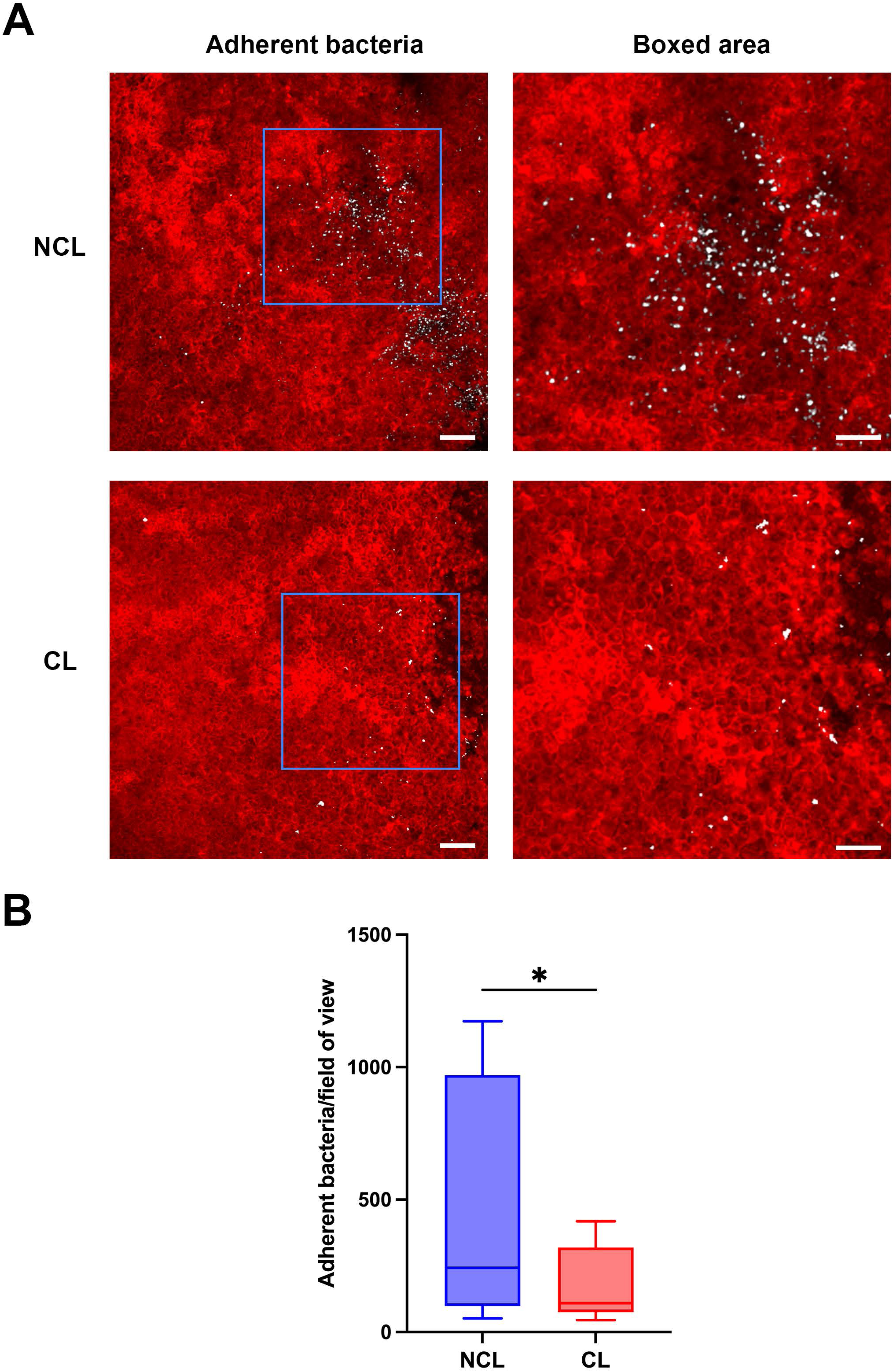
Prior contact lens wear is associated with reduced adherence of commensal bacteria to a superficially-injured mouse cornea. A) Images showing superficially-injured corneas of mT/mG-LysMcre mice (red cell membranes) with (CL) or without (NCL) prior contact lens wear (4-6 days). Adherent bacteria, *Macrococcus epidermidis* (white), were detected using a 16S rRNA-targeted FISH probe. Right panels show higher magnification of the blue boxed areas with adherent bacteria. Scale bars: left 50 μm, right 30 μm. B) Quantification of adherent bacteria per field of view, showing a significant reduction of the median bacterial adhesion with prior lens wear. *P < 0.05 (Wilcoxon signed-rank test).

**Fig. 2.**
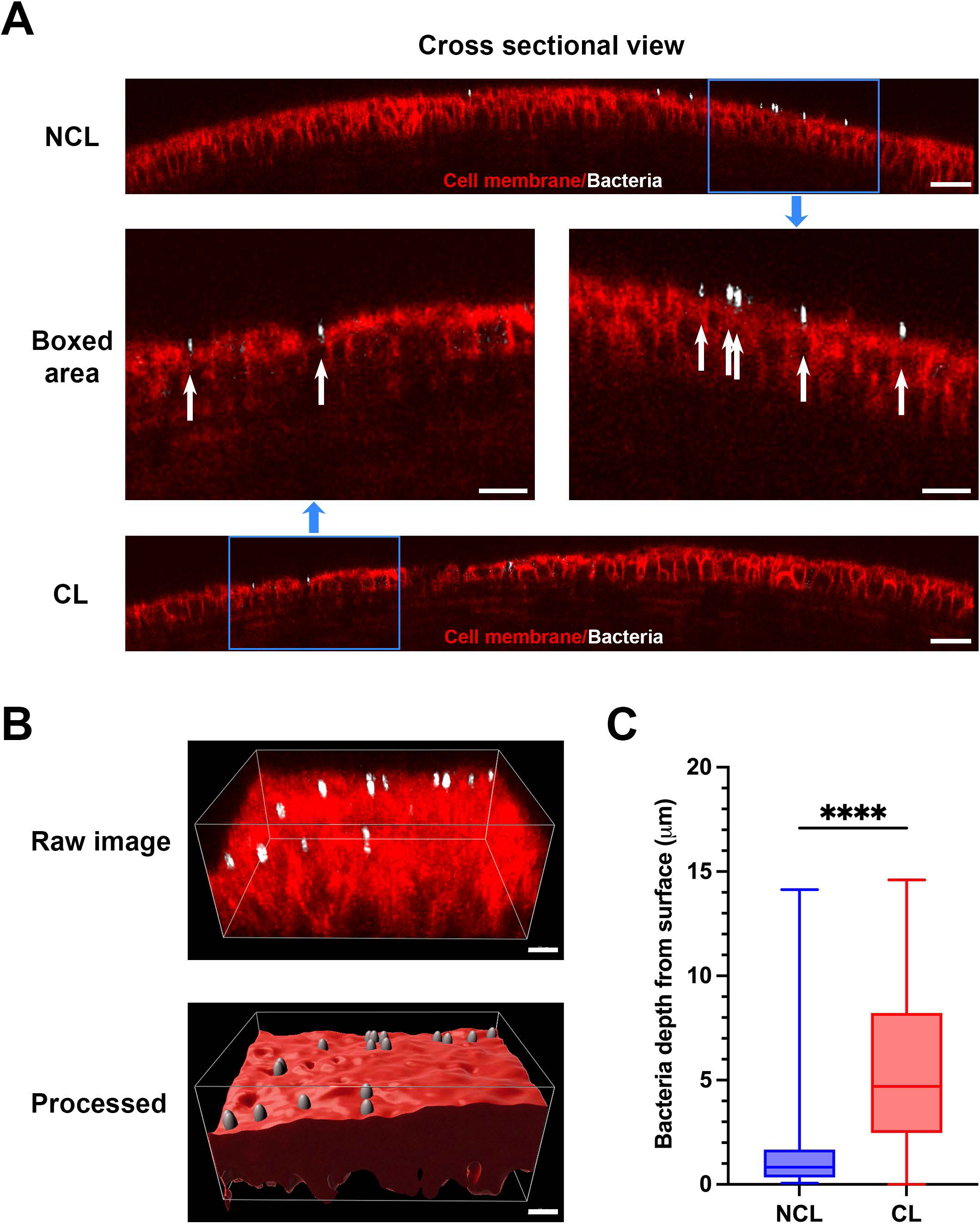
Prior contact lens wear is associated with deeper entry of the corneal epithelium by commensal bacteria adhering after superficial injury. A) Spatial positions of adherent bacteria (white) on the superficially-injured corneal epithelium (red) shown in cross-sectional view with (CL) and without (NCL) prior lens wear. Middle panels showed blue boxed areas from upper and lower panels in a higher magnification with bacteria indicated by white arrows. Scale bars: upper and lower 30 μm, middle 20 μm. B) Imaris spots rendering of adherent bacteria and surface rendering of the corneal epithelium with depth of penetration measured by distance of the spots to the surface. Scale bar 10 μm. C) Quantification of median bacterial depth from the epithelium surface showing deeper penetration with prior contact lens wear. ****P < 0.0001 (Mann-Whitney U test).

### Prior contact lens wear induced corneal Lyz2+ cell recruitment with cells moving closer to the epithelium surface

Since we previously showed Lyz2^+^ cell infiltration in lens wearing corneas after 6 or 7 days wear in the absence of deliberate bacterial inoculation or superficial injury (11), we compared Lyz2^+^ cell numbers and morphology in prior lens wear versus no lens wear corneas after superficial injury and inoculation with commensal bacteria. A ∼75 % increase in corneal Lyz2^+^ cells was observed in prior lens-wearing corneas vs. no lens wear contralateral eyes (Fig. 3A, B) (P = 0.047, Student’s t-test). More Lyz2^+^ cells were found in the peripheral vs. central cornea in both groups (Fig. 3A, B). Increased Lyz2^+^ cell numbers closely correlated with decreased adherence of commensal bacteria when analyzing central and peripheral corneas in both groups (Fig. 3C) (Pearson correlation coefficient r = - 0.9668, P = 0.033).

**Fig. 3.**
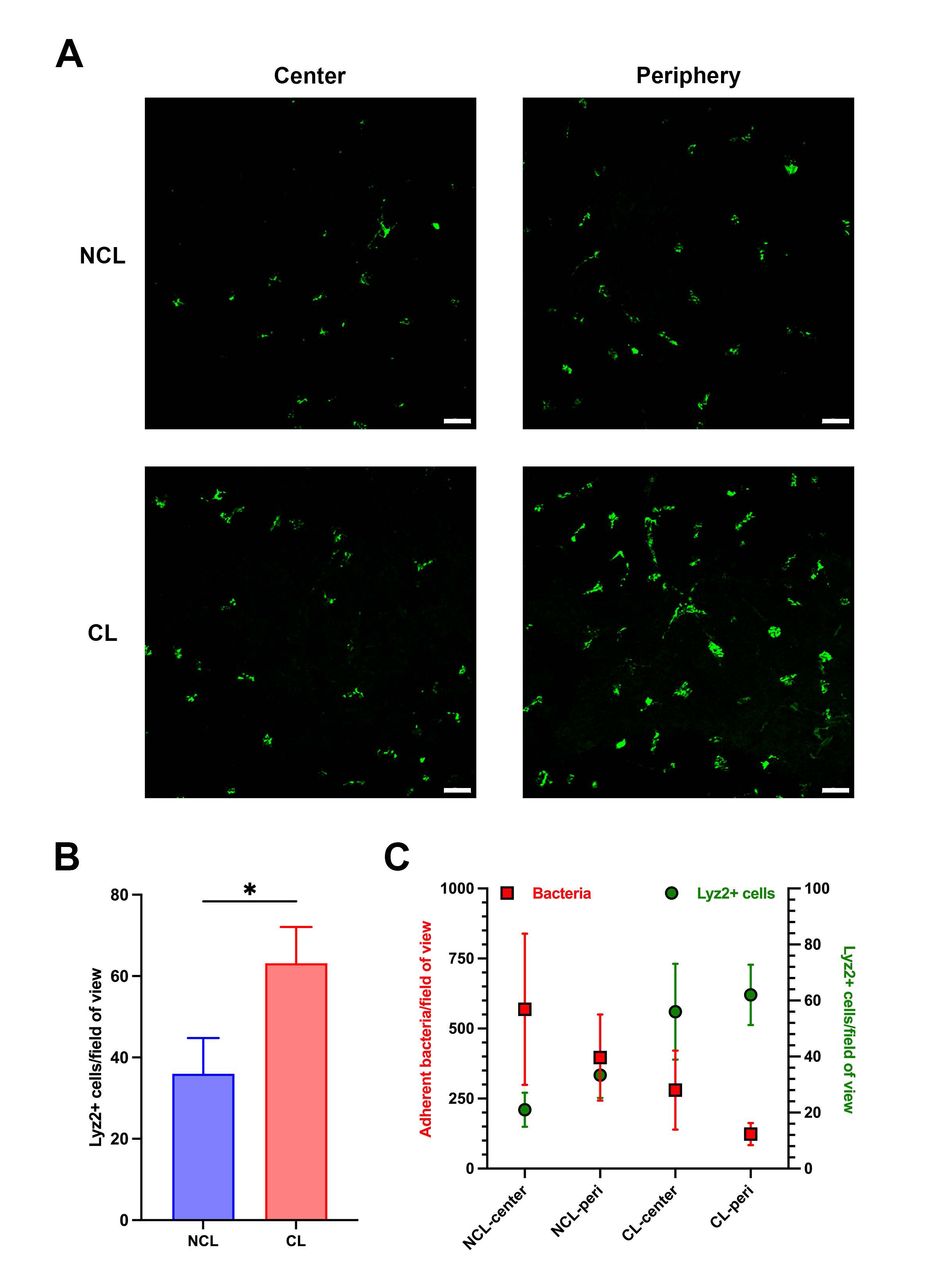
Prior contact lens wear is associated with greater Lyz2^+^ cell numbers in superficially-injured corneas after inoculation with commensal bacteria versus no lens wear. A) Maximum intensity z-projection of the GFP signal from Lyz2^+^ cells in the mT/mG-LysMcre mouse cornea after superficial-injury and commensal bacteria inoculation (4 hours). Lyz2^+^ cell distributions are shown in central and peripheral areas of the cornea with (CL) or without (NCL) lens wear (4-6 days). Scale bar 50 μm. B) Corneal Lyz2^+^ cell numbers per field of view with or without lens wear. *P < 0.05 (Student’s t-test). C) Correlation analysis shows a strong negative correlation between commensal bacteria adhesion and Lyz2^+^ cell numbers comparing central and peripheral corneas with and without prior lens wear (Pearson correlation coefficient r = - 0.967, P = 0.033).

Examination of Lyz2^+^ cell location in superficially-injured corneas after commensal bacteria inoculation showed that these cells were either within the epithelium or extending processes into the epithelium in both prior lens wear and no lens wear groups (Fig. 4A) with more cells showing this phenotype in prior lens-wearing corneas (Fig. 4B) (P = 0.031, Student’s t-test). Lyz2^+^ cells were also closer to the corneal surface in prior lens-wearing corneas versus no lens wear with these differences apparent only in the peripheral cornea (Fig. 5A). For example, Lyz2^+^ cell distance from the corneal endothelium in the peripheral cornea was: NCL 83.9 [77.2 – 90.0] vs. CL 93.8 [80.0 – 109.0] μm (P < 0.0001, Mann-Whitney U test). Analysis of Lyz2^+^ cell morphology showed a small increase in cell sphericity in prior lens-wear corneas versus no lens wear in peripheral cornea cells (Fig. 5B). For example, in the peripheral cornea with 1.0 being most spherical: NCL 0.70 [0.64 - 0.76] versus CL 0.71 [0.64 - 0.80] (P = 0.04, Mann-Whitney U test). However, overall changes in cell sphericity were not significant (Fig. 5B). No overall change in Lyz2^+^ cell volume was detected between prior lens-wear and no lens wear corneas after commensal inoculation (Fig. 5C), although opposing differences were observed if central and peripheral corneas were compared. For the central cornea, median cell volume increased with prior lens wear: NCL 1,053 [112 – 2,470] vs. CL 1,493 [440 – 3,026] μm^3^ (P = 0.001, Mann-Whitney U test). In contrast, peripheral cornea median cell volume decreased with prior lens wear: NCL 2,152 [1,357 – 3,046] vs. CL 1,844 [744 - 3,309] μm^3^ (P = 0.034, Mann-Whitney U test, Fig. 5C).

**Fig. 4.**
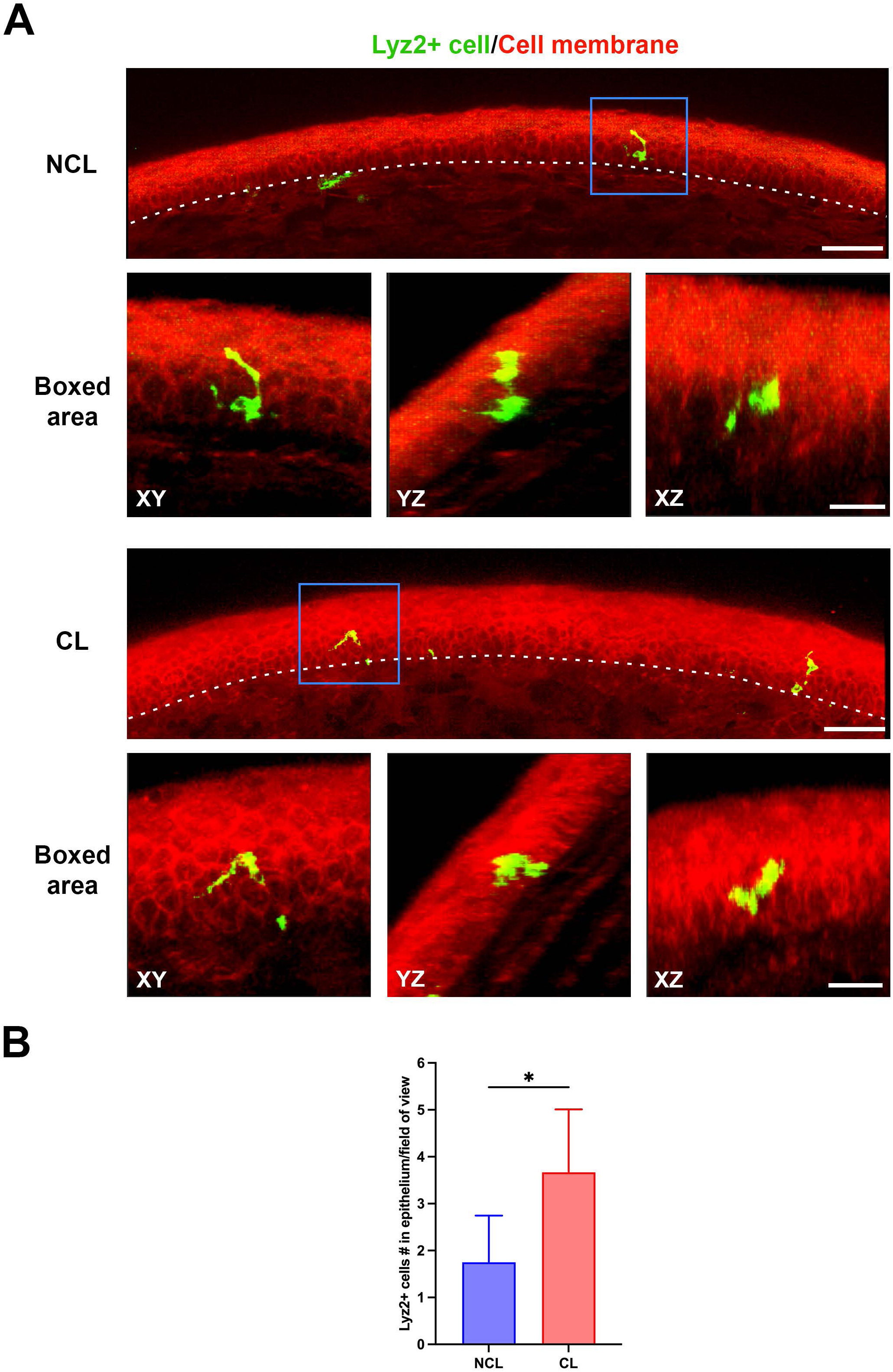
Prior lens wear is associated with more Lyz2^+^ cell migration into the corneal epithelium. A) Extended sectional view (15 μm) of a superficially-injured mT/mG**-**LysMcre mouse peripheral cornea after commensal bacteria inoculation (4 hours) showing Lyz2^+^ cells (green) inside or extended processes into the corneal epithelium (basal membrane indicated by white dotted lines) with (CL) and without (NCL) prior lens wear. Representative three-dimensional images of Lyz2^+^ cells in the corneal epithelium indicated by blue boxes and displayed in XY, YZ and XZ planes at higher magnification. Scale bar 50 μm or boxed area 20 μm. B) Quantification of Lyz2^+^ cell number per field of view showing increased Lyz2^+^ cells inside or extended processes into corneal epithelium with prior contact lens wear. *P < 0.05 (Student’s t-test).

**Fig. 5.**
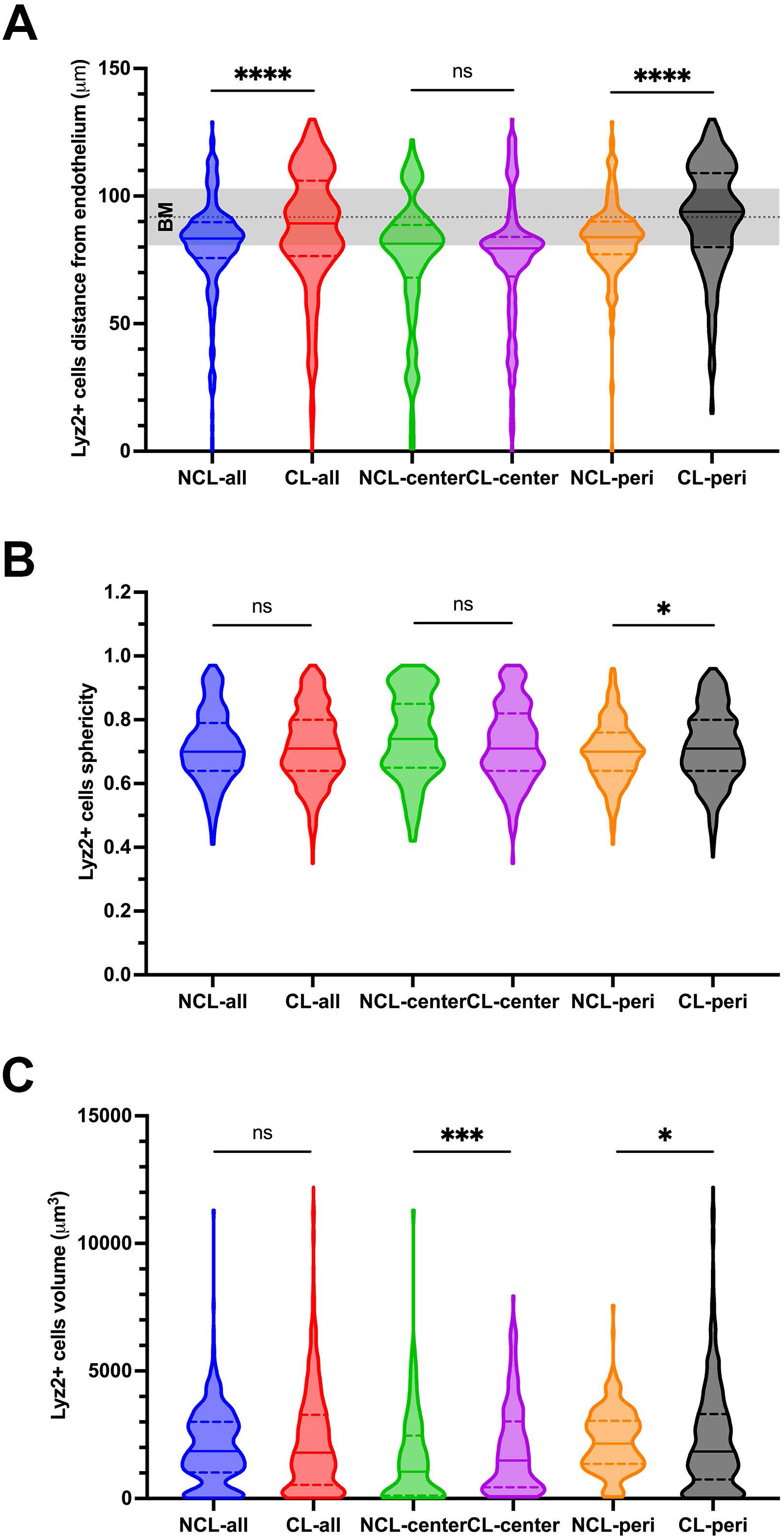
Distributional and morphological changes of Lyz2^+^ immune cells in the superficially- injured cornea after commensal bacteria inoculation. A) Lyz2^+^ cells showed increased migration towards the corneal epithelium through the basal membrane (BM) in prior lens wear cornea, specifically in the peripheral (peri) cornea. B) Lyz2^+^ cells showed little difference in sphericity with and without prior lens wear albeit with a statistically significant increase in the peripheral cornea with prior lens wear (1 = most spherical). C) Lyz2^+^ cell volume showed opposing changes with prior lens wear when comparing the central (increased) vs. peripheral (decreased) cornea versus no lens wear. *P < 0.05, ***P < 0.001, ****P < 0.0001, ns = not significant (Mann-Whitney U test).

### Prior lens wear reduces *P. aeruginosa* adhesion to the cornea and its epithelium penetration

We next tested the effect of prior lens wear on susceptibility of the superficially-injured cornea to the opportunistic pathogen *Pseudomonas aeruginosa*. As observed for commensal bacteria, fewer *P. aeruginosa* adhered to the cornea with prior contact lens wear vs. no lens wear, a ∼45 % decrease (P = 0.048, Student’s t test) (Fig. 6A, B). In contrast to the commensal, *P. aeruginosa* showed reduced epithelium penetration with prior lens wear, NCL 3.47 [2.13 - 5.43] vs. CL 2.50 [1.50 - 4.38] µm (P < 0.0001, Mann-Whitney U test) (Fig. 6C). Reduced *P. aeruginosa* adhesion with prior lens wear was also associated with increased Lyz2^+^ cell numbers in those corneas (∼2.9-fold) (P = 0.039, Student’s t-test) (Fig. 6D). However, in contrast to the commensal, the correlation analysis between reduced bacterial adhesion and increased Lyz2+ cell numbers for central and peripheral corneas for both groups was not significant (Fig. 6E) (Pearson correlation coefficient r = - 0.7736, P = 0.226).

**Fig. 6.**
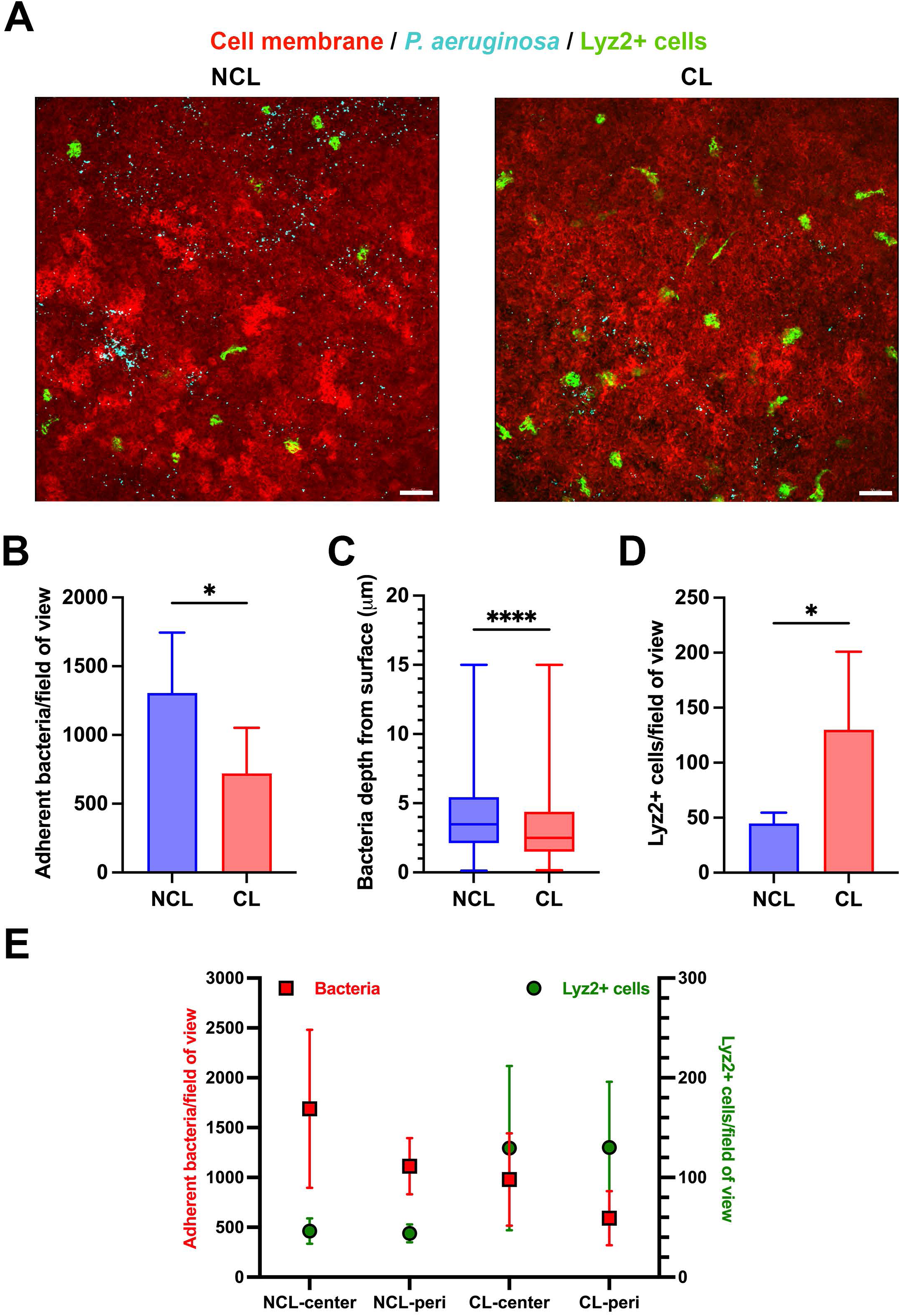
Prior contact lens wear reduces *P. aeruginosa* adhesion and epithelium penetration. A) Representative images show fewer adherent *P. aeruginosa* (Cyan) on the cornea of a mT/mG- LysMcre mouse after superficial-injury and bacterial inoculation (4 hours) with prior lens wear (4- 6 days) vs. no lens wear. Scale bar 50 μm. B) Quantification showing significantly reduced adherent *P. aeruginosa* per field of view with prior lens wear. C) Reduced penetration of *P. aeruginosa* into the corneal epithelium with prior lens wear, and D) Increased Lyz2^+^ cells per field of view with prior lens wear. For B and D, *P < 0.05 (Student’s t-test). For C, ****P < 0.0001 (Mann-Whitney U test). E) Correlation analysis shows a negative correlation between *P. aeruginosa* adhesion and Lyz2^+^ cell numbers comparing central and peripheral corneas with and without prior lens wear, but the P value was not significant (Pearson correlation coefficient r = - 0.7736, P = 0.2264).

Fig. 7 shows Lyz2^+^ cell distribution and morphology changes in the superficially-injured cornea after *P. aeruginosa* inoculation with and without prior lens wear. Most Lyz2^+^ cells were closer to the endothelium with prior lens wear: Overall NCL 82.4 [56.2 – 100.0] vs. CL 71.4 [47.5 - 91.7] µm from the endothelium (Fig. 7A) (P < 0.0001, Mann-Whitney U test). That difference reflected peripheral cornea Lyz2^+^ cells. Although a sub-group in the central cornea appeared to migrate towards the epithelium with prior lens wear (see Fig. 7A), that difference not significant vs. no lens wear. Lyz2^+^ cells were more spherical in prior lens wear corneas versus no lens wear, overall and in both the central and peripheral cornea: for example in the central cornea (NCL 0.81 [0.70-0.87] vs. CL 0.84 [0.75-0.89]) P < 0.001 (Mann-Whitney U test) (Fig. 7B). Lyz2^+^ cells were smaller in volume with prior lens wear but only in the central cornea (NCL 2,722 [1,331-3,891] vs. CL 1,896 [1,297-2,922] μm^3^ (P = 0.001, Mann-Whitney U test) (Fig. 7C).

**Fig. 7.**
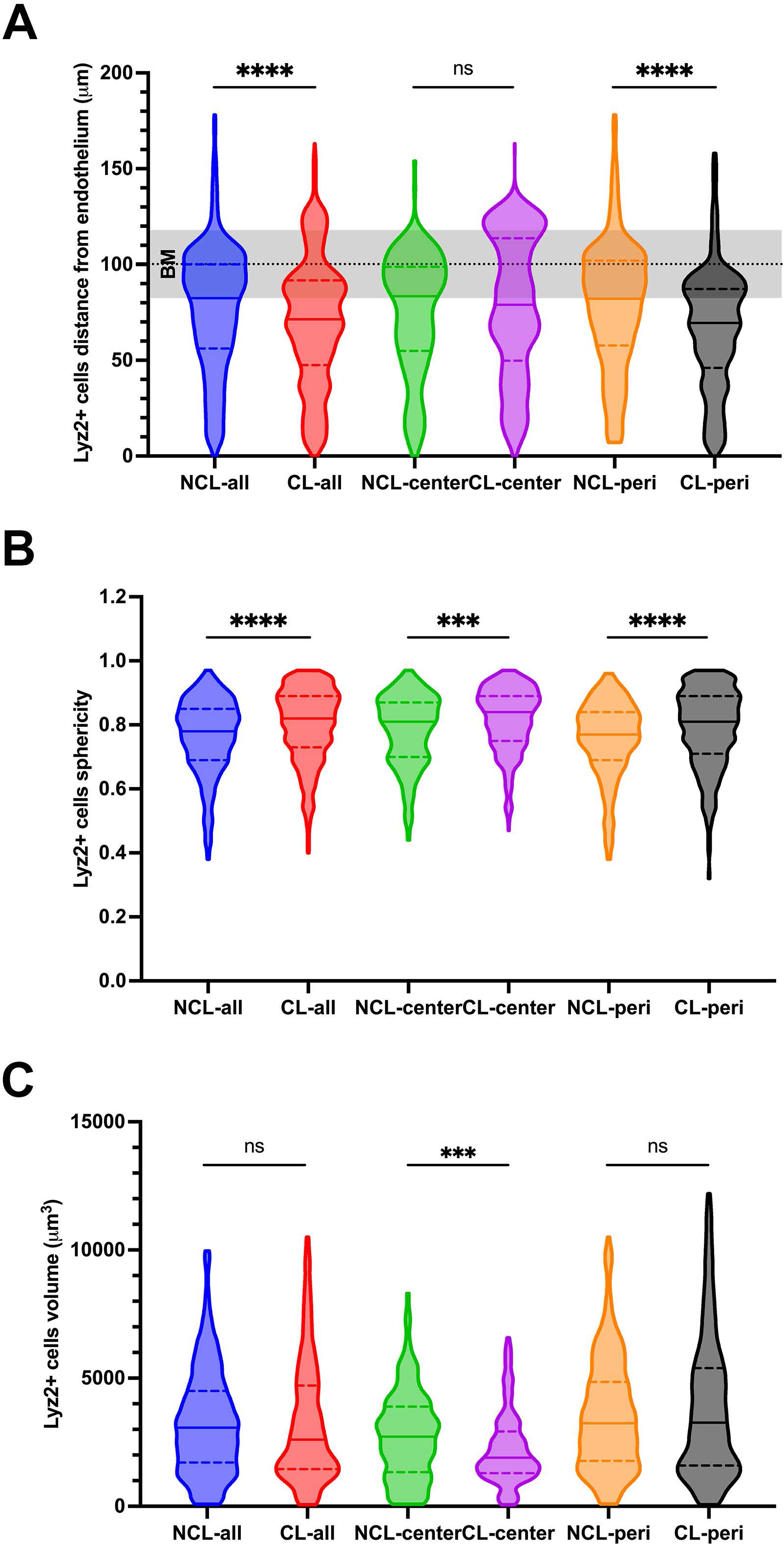
Distributional and morphological changes of Lyz2^+^ immune cells in a superficially- injured cornea after *P. aeruginosa* inoculation. A) Lyz2^+^ cells were further away from the epithelium with prior lens wear corneas but only in the peripheral cornea, and B) Lyz2^+^ cells were more spherical in prior lens wear corneas, and C) Lyz2+ cells showed a decrease in cell volume with prior lens wear. *P < 0.05, ***P < 0.001, ****P < 0.0001, ns = not significant (Mann-Whitney U test).

## Discussion

Here, we used a murine model of contact lens wear to study the impact of 4-6 days prior lens wear on corneal defenses against bacterial adhesion using both commensal and pathogenic bacteria. To enable bacterial adhesion to the otherwise resistant corneal surface, tissue paper blotting was used (22). The results showed that prior lens wear enabled a significant protective effect against bacterial adhesion of both the commensal and pathogenic bacteria, in both cases associated with increased Lyz2^+^ cell numbers. Thus, prior lens wear over 4-6 days has a protective effect against adhesion of two very different bacteria: a murine commensal (Gram-positive) and a versatile opportunistic pathogen (Gram-negative) and therefore the modification of one or more corneal defenses against bacterial adhesion over the prior lens wearing period.

While prior lens wear protected against bacterial adhesion for both types of bacteria, differences in the details were noted. For example, the adherent bacteria penetrated further into the epithelium for the commensal, while *P. aeruginosa* instead showed a reduced ability to penetrate. For the commensal, the Lyz2^+^ cells moved closer to the adherent bacteria and in the central cornea were larger compared to when a lens had not been worn, whereas for *P. aeruginosa* the Lyz2+ cells were further away and smaller in the central cornea. Their sphericity was increased by prior lens wear for *P. aeruginosa* with no overall change for the commensal. The differences in bacterial location and in the Lyz2^+^ cell response for the commensal versus *P. aeruginosa* challenge suggest different mechanisms might be at play depending on bacterial type.

It is still unclear whether alterations to the Lyz2^+^ cell responses to the injury/bacteria induced by prior lens wear are actually involved in protection against adhesion, or whether they are simply correlative. Lyz2^+^ cells include monocytes, macrophages, a subset of CD11c^+^ (dendritic cells), and neutrophils each of which can, or has the potential to, modulate the resistance of the cornea to bacterial adhesion. The role of neutrophils, monocytes and macrophages in responding to and defending already infected corneas against pathogens such as *P. aeruginosa* is well established (24–27).

More directly relevant to this study, we have used the same superficial injury/*P. aeruginosa* inoculation model without prior lens wear to study CD11c^+^ cell responses and found them migrating *<u>towards</u>* the *P. aeruginosa* adhering to the corneal surface. Depleting the CD11c^+^ cellsfurther enhanced bacterial adhesion (28) showing that CD11c^+^ cells can protect superficially- injured corneas against *P. aeruginosa* adhesion. Here, we introduced prior lens wear to that superficial injury/*P. aeruginosa* challenge model and found that the Lyz2^+^ cells (a subset of which are CD11c^+^ cells) were now *<u>further away</u>* from the adherent bacteria compared to no prior lens wear despite prior lens wear protecting against *P. aeruginosa* adhesion. This suggests that the role of immune cells in protecting superficially-injured corneas against *P. aeruginosa* is likely to be complex and might even depend upon circumstances. Previously, we showed that Ly6G^+^ cells (likely neutrophils) infiltrate the cornea 5-6 days into uninoculated lens wear in this murine contact lens model (11). Thus, neutrophils (also Lyz2^+^) are another potential candidate for involvement in countering bacterial adhesion after prior lens wear. Other immune cells (e.g. γδ T cells) involved in corneal parainflammatory responses to uninoculated lens wear (11,14–16) may also contribute to the defenses against bacterial adhesion after lens removal shown here. At present, however, a direct connection to lens-induced corneal parainflammation remains to be shown.

It is also possible that the Lyz2^+^ cell changes noted are simply correlative and that other factors, possibly beyond immune cells, are responsible for reducing bacterial adhesion associated with prior lens wear. Various non-immune cell factors have been shown by us, and others, to counter bacterial adhesion at the corneal surface. They include corneal mucins, lectins, antimicrobial peptides (including defensins, keratin derived antimicrobial peptides), DMBT1, surfactant proteins, epithelial cell polarity, and junction-related factors (29–36). Moreover, we recently reported that corneal defenses against adhesion include their sensory nerves, with TRPA1 involved in countering *P. aeruginosa* (37), and TRPV1 countering both environmental bacteria and *Staphylococcus aureus* (37,38). Moreover, we have shown that these receptors are also required for contact lens-induced corneal parainflammation and modulation of resident immune cells in the murine lens wear model (14). This raises the possibility that sensory nerve-associated TRP receptors are also involved in how prior lens wear reduces the propensity for bacterial adhesion. Ultimately, the defense mechanisms involved are likely to be multifactorial possibly involving a combination of sensory nerves, immune cells and changes to intrinsic anti-adhesive or directly antimicrobial defenses. Additional studies will be needed to tease apart the mechanism(s) by which prior lens wear reduces bacterial adhesion to the cornea.

How does prior lens wear protecting the cornea against *P. aeruginosa* adhesion align with the fact that contact lens wear predisposes the cornea to *P. aeruginosa* infection? Here, the lens was deliberately removed before the bacteria were inoculated into the eye. This was to allow us to tease apart the contribution of lens wear itself to defenses against adhesion, the hypothesis being that it induces a protective response based on our observation that there is parainflammation and its significance remained unknown. Removing the lens before adding the bacteria is different from keeping the lens in place whilst the bacteria are present for several reasons. Firstly, contact lens-associated corneal infection in animal models requires a lens deliberately inoculated with *P. aeruginosa* to remain in place on the cornea for up to 13 days in the mouse (11) or for 7-10 days in the rat (10). Without a lens, *P. aeruginosa* and other bacteria are rapidly cleared from the ocular surface (20,21). Their inability to gain a foothold on the intact healthy adhesion resistant cornea is because “antimicrobial factors” present at the ocular surface/tear fluid can kill them, incapacitate them, aggregate them, repel them and/or manipulate their gene expression (20,21,29,34,35,39–43). Secondly, we have shown that the posterior surface of lenses in *P. aeruginosa* infected eyes of rats harbored *P. aeruginosa* biofilms, a sign of bacterial adaptation to the ocular surface environment (10). This adaptation to the ocular surface on a lens (i.e. exposure to both simultaneously) appears to be a key feature in the pathogenesis of contact lens-related infection, as transferring colonized lenses from rat eyes with *P. aeruginosa* infections into eyes of naïve rats enabled infections to develop more quickly (∼2 days) (10). Additionally, we have shown that if *P. aeruginosa* is exposed to tear fluid for several hours (not normally feasible without lens wear due to clearance mechanisms), the bacteria release outer membrane vesicles able to kill epithelial cells at the surface of the cornea, and that this in turn promotes corneal epithelium susceptibility to *P. aeruginosa* adhesion (44). Also relevant to this discussion, we previously showed that superficial injury to the cornea alone does not enable infection susceptibility despite it enabling bacterial adhesion (22). This is because there are deeper layer defenses in place that prevent adherent bacteria from progressing further and superficial injury alone is insufficient to compromise them (28). Thus, there are many factors contributing to the pathogenesis of contact lens-related corneal infection beyond bacterial adhesion, including changes to both the bacteria and host while a lens remains *in situ* (45). As such, results of this study showing reduced bacterial adhesion to corneas that had previously worn a lens adds a small piece to the complex puzzle of how the cornea adapts to lens wear and responds to microbes, information that will help to eventually understand and prevent pathogenesis of contact lens related and other infections.

In conclusion, contact lens wear in humans causes both asymptomatic parainflammation and acute corneal infection. While animal models of lens wear can reproduce both events, the relationship between them remains unclear. The present study adds to our knowledge by showing that 4-6 days of lens wear in mice, which enables parainflammation, can render the cornea less susceptible to adhesion by either a commensal or pathogenic *P. aeruginosa* if inoculated after the lens is removed. This differs from events occurring if the lens is left in place, which instead leads to infection. Whether and how the increased corneal Lyz2^+^ immune cell numbers and changes to their location and morphology are involved in the protective phenotype for each bacterial type remains to be determined. Follow up studies will also be needed to understand how continuing lens wear during *P. aeruginosa* challenge undermines this defensive response to lead to infection.

## Acknowledgements

Our thanks to Dr. Alain Filloux (Imperial College London, UK) for originally providing the *P. aeruginosa* wild-type strain PAO1F.

